# Right Temporoparietal Junction Encodes Mental Experience of Others More Than Evaluating False Versus True Belief

**DOI:** 10.1101/2021.06.10.447930

**Authors:** Branden J. Bio, Arvid Guterstam, Mark Pinsk, Andrew I. Wilterson, Michael S. A. Graziano

**Affiliations:** Department of Psychology, Princeton University, Princeton, NJ 08544; Princeton Neuroscience Institute, Princeton University, Princeton, NJ 08544

**Keywords:** Theory of Mind, Mentalizing, Temporoparietal Junction, False belief task, fMRI

## Abstract

When people make inferences about other people’s minds, called theory of mind (ToM), a network in the cerebral cortex becomes active. ToM experiments sometimes use the false belief task, in which subjects decide whether a story character believes A or B. The “false” belief occurs if the character believes A when B is true. We devised a version in which subjects judged whether a cartoon head “believed” a ball to be in box 1 or box 2. The task was a visual, reaction-time version of a ToM task. We proposed two alternative hypotheses. In hypothesis 1, cortical regions of interest within the ToM network should distinguish between false and true belief trials, reflecting outside information that the subjects have about the cartoon character. In hypothesis 2, the ToM network should distinguish between conditions only if the subjects think that the cartoon character can distinguish between the conditions, thus reflecting a model of the internal contents of the cartoon character’s mind. The results supported hypothesis 2. Events that the cartoon could not “see” did not affect activity in the ToM network; the same events, when the cartoon could apparently “see” them, significantly affected activity in the right temporoparietal junction (TPJ). The results support the view that the right TPJ participates in modeling the mental states of others, rather than in evaluating the accuracy of the beliefs of others, and may help explain why previous experiments showed mixed results when directly comparing false belief to true belief conditions.

**Significance statement:** How do we understand the minds of others? A specific network of areas in the human brain is known to emphasize building models of other people’s minds. In one traditional hypothesis, this network helps us distinguish whether someone else’s beliefs are true or false. Here we show that the network is relatively insensitive to one’s own, outside knowledge about whether someone else’s beliefs are true or false. Instead, at least one node in the network appears to limit itself mainly to reconstructing the other person’s point of view. This social cognition network may be more about empathy than about critical evaluation.

**Classification:** Social Sciences - Psychological and Cognitive Sciences

## Introduction

Building a model of other people’s thoughts, emotions, and beliefs, also called theory of mind (ToM), is foundational to our social lives [1–4]. A large literature shows that ToM tasks tend to activate a specific network of areas in the human cerebral cortex [5–19]. Those areas typically include the temporoparietal junction (TPJ), sometimes bilaterally but with a bias toward the right side; the superior temporal sulcus (STS), again sometimes bilaterally but with a bias toward the right side; the medial prefrontal cortex (MPFC); and the precuneus. Other brain areas are also reported in ToM studies, such as the temporal pole and the amygdala, but the areas listed above are often more consistently active during ToM tasks, as shown in meta-analysis studies [12,16,18].

Experiments on the ToM cortical network often use a classic paradigm called the false belief task [4,10]. In it, the subject of the experiment must decide, based on information known to be available to a character in a story, whether the character thinks that A or B is true. For example, if Sally originally put her sandwich into box A, will she still think it is in box A, even after someone else, unbeknownst to Sally, has moved it to box B? To answer correctly, the subject of the experiment must have a sophisticated enough theory of Sally’s mind to realize that Sally can believe something that is false – that is distinct from the world around her – depending on the information available to her.

In a recent behavioral study, we designed a variant of the false belief task [21]. Our version was designed to build up a set of information visually over the course of each trial. Visual ToM tasks incorporating cartoons, or, more often, videos, have been used before [8,13,22–26]. As shown in Figure 1A, first, two cartoon heads appeared, looking at two open boxes. Second, the “ball,” a red dot, appeared in one box. Third, one head was shown having its eyes covered such that it could no longer see what was in either box, while the other head remained with uncovered eyes. Fourth, on half of the trials, the ball switched from one box to the other. Fifth and finally, a question mark appeared in one head or the other. The task was to decide whether the head indicated by the question mark “believed” the ball to be in box 1 or box 2. The purpose of this incremental presentation of information, and the use of two heads, was to ensure that the subjects would not know how to answer the question until the last pictorial piece of information, the question mark, was presented. At that moment, subjects had sufficient information to make a ToM judgment, and they responded in a speeded manner within a limited response window (1.5 s). The design therefore converted the false belief task into a pictorial, reaction-time task, in which a single event (the appearance of the question mark) triggered the moment when subjects needed to make a ToM judgment. With all stimuli being fully right-versus-left counterbalanced across trials, the conditions were nearly visually identical, providing a good control for each other.

**Figure 1.**
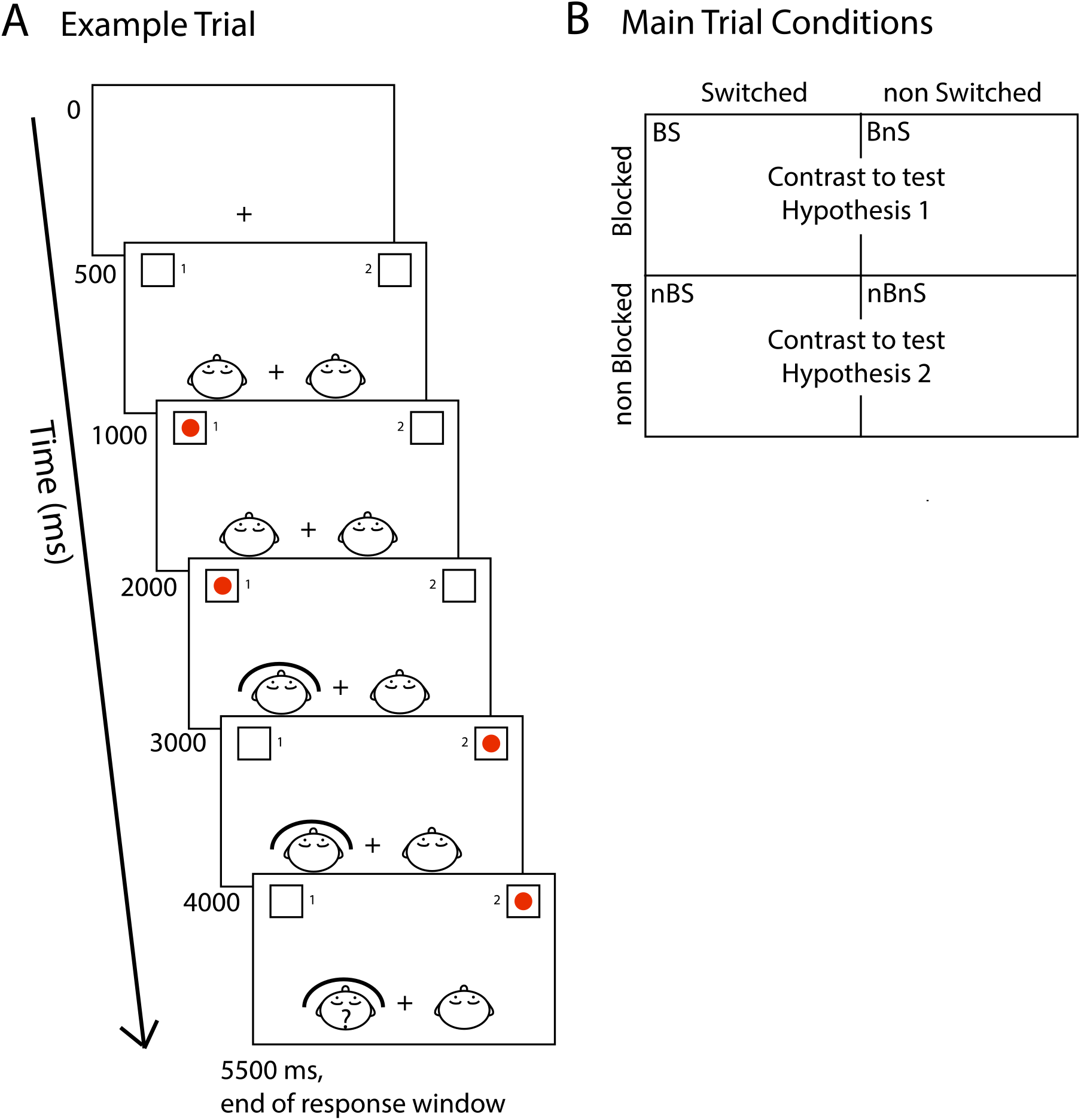
Behavioral paradigm. A. Timeline of events during a typical trial. Fixation point appeared at start of trial. Then two heads and two boxes appeared. Then a red ball appeared in one box. Then one head had its sight blocked by the curved barricade. Then, on half of trials, the ball switched to the opposite box. Then a question mark appeared in one head, signaling subjects to respond by deciding whether the indicated head “thinks” the ball is in box 1 or box 2. All events were right-left counterbalanced among trials. B. Four main trial conditions formed by the 2 × 2 design of blocked versus nonblocked configurations X switched versus nonswitched configurations. BS and BnS trials were contrasted to test hypothesis 1, whereas nBS and nBnS trials were contrasted to test hypothesis 2.

We conceptualized the paradigm as a 2×2 design, as shown in Figure 1B. The first variable was whether the indicated cartoon head had its vision blocked: in half the trials, the subjects had to judge the visual beliefs of the cartoon head whose eyes were covered, whereas in the other half of trials, the subjects had to judge the visual beliefs of the cartoon head whose eyes were uncovered. The second variable was whether the ball switched from one box to the other: in half the trials, the ball switched boxes midway through the trial, whereas in the other half of trials, the ball began in one box and remained there without switching. This design resulted in four trial conditions that we termed blocked-switched (BS), blocked-nonswitched (BnS), nonblocked-switched (nBS), and nonblocked-nonswitched (nBnS).

In the present experiment, subjects performed this pictorial ToM task in a magnetic resonance imaging (MRI) scanner, which measured brain activity evoked by the subjects’ ToM decisions on each trial. The experiment differed from many previous brain imaging ToM experiments in two major ways. First, many previous experiments used a block design, relying on an average of brain activity across many seconds as subjects performed a continuous task such as reading a story [5,8,9,15,19,27]. In the present experiment, we analyzed brain activity evoked by a decision that occurred within a brief, reaction-time window. This focus on a narrow decision window was expected to result in a weaker brain activity signal, since it was averaged over a smaller time interval. We hoped, however, that focusing on a distinct moment of decision might increase the specificity of the results by reducing the influence of other aspects of the trial that were the same across conditions.

Second, many previous ToM experiments tested a contrast between a ToM condition, such as a false belief condition, and a nonsocial control, such as a false photo, false sign, or physical description condition [5,8,9,13,15,19,27]. Such designs are especially effective at revealing the cortical ToM network. However, many incidental, unintended differences might be present between social and nonsocial tasks, potentially complicating the interpretation of results. Here, we examined differences among ToM conditions that were visually and cognitively nearly identical, but that involved slightly different ToM decision processes at the crucial moment during the trial. We hoped that this design would help test more specific hypotheses about the cortical ToM network.

Although many different comparisons are possible among the four conditions (and are shown in the **Supplementary Information**), here we focus on two specific hypotheses. In hypothesis 1, brain areas in the ToM network should respond differently to false belief and true belief trials. Many previous studies have compared false belief to true belief conditions, to test whether false belief reasoning requires especially complex or intensive ToM, or is processed in a specific part of the ToM network as distinct from true belief reasoning [23,25,28,29]. The results of these previous studies are mixed. Though there is evidence of false belief processing emphasized in some subregions of the ToM network, especially in the TPJ and the MPFC [25,28,29], some researchers have argued that false and true belief conditions are similar in how they recruit ToM reasoning [15]. Here, to help address this ongoing question, we tested whether false belief trials and true belief trials affect ToM brain areas differently. The BS trials correspond to the only false belief condition in our task. The head sees the ball in one box, then the head’s eyes are covered, then the ball is switched to the other box. To answer correctly, subjects must realize that the head “believes” the ball to be in the first box, even though the subjects can see ball is actually in the second box. In contrast, the BnS trials represent a balanced true-belief condition. In the BnS trials, once again the head sees the ball in one box and then the head’s eyes are covered. The ball, however, is not switched to the opposite box, and therefore, by the end of the trial, the head should believe the ball to be in the box in which it actually resides. The head has a “true” belief. The two trial types are closely matched except that one is a false belief condition and the other is a true belief condition. If hypothesis 1 is correct, then at least some cortical areas known to be involved in ToM should show significant activity in the comparison between BS and BnS trial types, indicating that they process the difference between false and true belief in others.

We proposed hypothesis 2 as an alternative to hypothesis 1. In hypothesis 2, the ToM brain areas build models of the cartoon character’s mind, regardless of whether the character’s beliefs are true or false. The crucial factor affecting the activity in the ToM brain areas, therefore, is not broadly what the subject knows, but only narrowly what the subject thinks the cartoon character knows. If the cartoon character is supposed to know the same information in two conditions – i.e. have the same mental experience – then the ToM brain areas should not distinguish between those conditions. If the cartoon character is supposed to know different information in two conditions – i.e. have different mental experiences – then the ToM brain areas should react differently to those conditions. Hypothesis 2 predicts a significant difference in brain activity, not between the BS and BnS conditions (as in hypothesis 1), but between the nBS and the nBnS conditions. To understand why, consider the following. In the BS and the BnS trials (the false belief and true belief conditions contrasted to test hypothesis 1), the cartoon head is supposed to have the same mental experience. The configuration, in both trial types, implies that the head “sees” the ball in a box, the head’s eyes are covered, and the head “believes” that the ball is in the same box in which it most recently saw it to be. In the perspective of the head, the two conditions are identical, and therefore the mental state that must be attributed to the head should be identical. Brain areas involved in reconstructing the mental states of others might therefore react in the same way to both conditions. The subject may know that the character has a false belief in one condition (BS) and a true belief in the other condition (BnS), but the cartoon character should not know that. Related to this hypothesis, it has been suggested [30] that parts of the ToM network, especially the right TPJ, are involved in filtering out or ignoring what is actually happening right now in one’s own experience and instead building a model of what could, hypothetically, be happening in another mind.

Now consider the difference between the nBS and nBnS trials. The difference here is that the cartoon’s eyes are not blocked, and therefore the character can see the ball at all times. In the nBS trials, the cartoon character can see the ball placed in one box, and can see the ball switched to the other box. By implication, the head might be thinking something like, “I first saw the ball there, then it switched, and now I know it’s currently here instead.” In contrast, in the nBnS trials, the cartoon character can see the ball placed in a box, and can see that it remains in that box. By implication, the head might think something like, “The ball is right here, where I see it to be.” Hypothesis 2 predicts that the contrast between nBS and nBnS trials should result in significant activity in the ToM network, because subjects are imputing different mental experiences to the cartoon head.

To summarize, hypothesis 1 suggests that the crucial factor affecting ToM brain areas is the subject’s own mental evaluation of what the cartoon character knows (does the cartoon character have a false belief or true belief?). If hypothesis 1 is correct, the ToM brain areas should respond differently in BS and BnS trials. In contrast, hypothesis 2 suggests that the crucial factor affecting ToM brain areas is the mind state that is attributed to the cartoon character. Regardless of what the subject may know, the ToM brain areas process what the cartoon character is inferred to know. If hypothesis 2 is correct, then the ToM brain areas should be insensitive to the BS versus BnS comparison, and instead respond differently in the nBS and nBnS trials. (See Figure 1B for a diagram of conditions and comparisons.)

To carry out the comparisons, we used multi-voxel pattern analysis (MVPA). The method trains a support-vector machine (SVM) to decode, on the basis of the MRI signal, whether one or another condition was presented on any given trial. If decoding is significantly above chance for a cluster of voxels in the brain, then that cluster contains information that differentiates the two conditions of interest. MVPA is considered to be more sensitive to the information content in voxels than more traditional subtraction methods. We used two MVPA approaches. First, for the main analysis, we used a targeted approach, testing specific ToM brain areas. We defined six regions of interest (ROIs) based on a previous meta-analysis of brain activity evoked during ToM tasks [18], in accordance with generally accepted guidelines in ROI analysis [31]. A similar MVPA method applied to the same, predefined, six ROIs has been used in a previous study [32]. The advantage of this method is that one does not need to perform a large battery of social cognition tasks as localizers to define the most common ToM network areas; instead one can rely on the extensive past literature to define these most common areas. The six areas based on prior meta-analysis [18] included the TPJ bilaterally, STS bilaterally, precuneus, and MPFC (see **Methods** for definitions of ROIs). Second, we used a searchlight analysis that tested the whole brain for clusters of significant decoding, statistically corrected for multiple comparisons across the brain. The searchlight analysis is less statistically sensitive and was used here as an exploratory method to test whether any areas of the brain might have shown especially strong decoding outside the ROIs.

## Methods

### Subjects

All subjects provided informed consent and all procedures were approved by the Princeton Institutional Review Board. We tested 28 healthy human volunteers (17 females, 27 right handed, aged 18-50, normal or corrected to normal vision). Subjects were recruited from a paid subject pool, receiving 40 USD for participation.

### Experimental setup

Before scanning, all participants received task instructions and completed practice trials on a laptop computer outside of the MRI scanner. During scanning, subjects laid in a supine position on the MRI bed and used an angled mirror mounted on top of the head coil to view a screen approximately 80 cm from the eyes, on which visual stimuli were projected using a digital light processing projector (Hyperion MRI Digital Projection System, Psychology Software Tools, Sharpsburg, PA, USA) with a resolution of 1920 × 1080 pixels at 60 Hz. A PC running MATLAB (MathWorks, Natick, MA, USA) and the Psychophysics Toolbox [33] were used to present visual stimuli. A 5-button response unit (Psychology Software Tools Celeritas, Sharpsburg, PA, USA) was strapped to the subjects’ dominant hand. Subjects used only the index and middle fingers to indicate responses.

### Behavioral task

The task events are illustrated in Figure 1A. Participants saw a cartoon that included two heads, two boxes, and a ball. The ball was located in one of two boxes and the participant had to decide whether a cartoon head would most likely believe the ball to be in box 1, to the left, or box 2, to the right. Participants responded by button press only at the end of the trial when one of the two cartoon heads was indicated as the target for the ToM judgment.

Each trial began with a black fixation cross at the center of a white background. Participants were instructed to fixate on the cross. After 500 ms, the fixation cross was joined by a top-down view of two cartoon heads and two numbered boxes. The heads were centered 3.25 degrees to the left (head 1) and right (head 2) of the vertical midline of the screen, positioned on the horizontal midline (at the same height as the fixation cross). The boxes were centered 12 degrees to the left (box 1) and right (box 2) of the midline, and 9 degrees above the horizontal midline. After another 500 ms, a red ball appeared in one of the two boxes (half of the trials in box 1, half of the trials in box 2). Participants had been told in the instruction period that, in this configuration, both heads could see where the ball was located. After 1000 ms, one of the heads was blocked with a curved partition directly in front of it (half of the trials blocking head 1, half of the trials blocking head 2). Participants had been told in the instruction period that the blocked head could no longer see either the boxes or the ball, but that the other head could still see everything as before.

In half of the trials, 1000 ms after the blocking partition appeared, the ball switched position to the opposite box. If it was initially in box 1, it moved to box 2; if it was initially in box 2, it moved to box 1. The head that was blocked should therefore “believe” the ball to be still in the original box, and the head that was unblocked should “see” the ball move to the new box. In the other half of trials, the ball did not switch positions.

Finally, 4000 ms after the start of the trial, a question mark appeared inside one of the heads (half of trials in head 1, half of trials in head 2). The question mark indicated which head was to be the target of the participant’s judgment. The participant was instructed to respond as quickly as possible once the question mark appeared. By pressing one of two buttons on the button box, the participant reported whether the indicated head would most likely think the ball was in box 1 or box 2. Participants were allowed a response window of 1500 ms. Trials on which participants exceeded the given time to respond were not included in the analysis. Participants responded within the correct time window on most trials (98%). After the response window, the display of heads and boxes disappeared and a variable, 1000 - 3000 ms inter-trial interval followed, after which the next trial began with the onset of the fixation cross.

In summary, the task included the following conditions: the red dot could be initially presented in box 1 or box 2; the blocking screen could be placed in front of the left or right head; the red dot could be switched to the opposite box or remain in the same box; and the question mark could be presented in the left or right head. This 2×2×2×2 design resulted in 16 trial types, presented in a counterbalanced and randomized order. The trial types were collapsed into four main conditions for purposes of analysis (see Figure 1B). These conditions formed a 2 × 2 design as follows: blocked trials, on which the head indicated by the question mark was blocked by the screen, versus nonblocked trials, on which the indicated head was not blocked by the screen; and switched trials, on which the ball moved to the opposite box, versus nonswitched trials, on which the ball remained in the initial box. As shown in Figure 1B, these four conditions were labeled as blocked switched (BS), blocked nonswitched (BnS), nonblocked switched (nBS), and nonblocked nonswitched (nBnS). Only the BS trials corresponded to a false belief condition, in which the indicated head should “believe” the ball to be in one box when it was actually in the opposite box. The other three conditions corresponded to true-belief trials, in which the indicated head should think the ball was in the box that it actually was in.

Participants performed 256 trials (64 per main condition), in 8 runs of 32 trials each. Each run took approximately 5.5 minutes to complete and included 5 s of baseline before the onset of the first trial and 10 s of baseline after the offset of the last trial.

### fMRI data acquisition

Functional imaging data were collected using a 3T MAGNETOM Skyra (Siemens Healthineers AG, Erlangen, Germany) scanner equipped with a 64-channel head/neck coil. Gradient-echo T2*-weighted echo-planar images (EPI) with blood-oxygen dependent (BOLD) contrast were used as an index of brain activity [34]. Functional image volumes were composed of 46 near-axial slices with a thickness of 3.0 mm (with no interslice gap), which ensured that the entire brain excluding the cerebellum was within the field-of-view in all subjects (80 × 80 matrix, 2.5 mm x 2.5 mm in-plane resolution, TE = 30 ms, flip angle = 75°). Simultaneous multi-slice (SMS) imaging was used (SMS factor = 2). One complete volume was collected every 1.5 s (TR = 1500 ms). A total of 1300 functional volumes were collected for each participant, divided into 8 runs (130 volumes per run). The first five volumes of each run were discarded to account for non-steady-state magnetization. A high-resolution structural image was acquired for each participant at the end of the experiment (3D MPRAGE sequence, voxel size = 1 mm isotropic, FOV = 256 mm, 176 slices, TR = 2300 ms, TE = 2.96 ms, TI = 1000 ms, flip angle = 9°, iPAT GRAPPA = 2). At the end of each scanning session, matching spin echo EPI pairs were acquired with reversed phase-encode blips, resulting in pairs of images with distortions going in opposite directions for blip-up/blip-down susceptibility-derived distortion correction.

### FMRI preprocessing

Results included in this manuscript come from preprocessing performed using FMRIPREP version 1.2.3 [35] (RRID:SCR_016216), a Nipype [36] (RRID:SCR_002502) based tool. Each T1w (T1-weighted) volume was corrected for INU (intensity nonuniformity) using N4BiasFieldCorrection v2.1.0 [37] and skull-stripped using antsBrainExtraction.sh v2.1.0 (using the OASIS template). Spatial normalization to the ICBM 152 Nonlinear Asymmetrical template version 2009c [38] (RRID:SCR_008796) was performed through nonlinear registration with the antsRegistration tool of ANTs v2.1.0 [39] (RRID:SCR_004757), using brain-extracted versions of both T1w volume and template. Brain tissue segmentation of cerebrospinal fluid (CSF), white-matter (WM) and gray-matter (GM) was performed on the brain-extracted T1w using fast [40] (FSL v5.0.9, RRID:SCR_002823).

Functional data was slice time corrected using 3dTshift from AFNI v16.2.07 [41] (RRID:SCR_005927) and motion corrected using mcflirt (FSL v5.0.9) [42]. This procedure was followed by co-registration to the corresponding T1w using boundary-based registration [43] with six degrees of freedom, using flirt (FSL). Motion correcting transformations, BOLD-to-T1w transformation and T1w-to-template Montreal Neurological Institute (MNI) warp were concatenated and applied in a single step using antsApplyTransforms (ANTs v2.1.0) using Lanczos interpolation.

Many internal operations of FMRIPREP use Nilearn [44] (RRID:SCR_001362]) principally within the BOLD-processing workflow. For more details of the pipeline see https://fmriprep.readthedocs.io/en/latest/workflows.html.

### MVPA analysis

We analyzed the data using MVPA, which tests whether patterns of brain activity can be used to decode the distinction between two conditions. It is a more sensitive analysis than the more common, simple univariate subtraction methods, and the study was designed from the outset to use MVPA (thus many trials per condition were included). Two independent MVPA comparisons were performed: BS versus BnS trials, and nBS versus nBnS trials. We tested both comparisons within a set of six ROIs.

The ROIs were defined as spheres centered on the statistical peaks reported in an activation likelihood estimation meta-analysis of 16 fMRI studies (including 291 subjects) involving ToM reasoning [18], in accordance with the approach used in Guterstam et al., 2021 [32] and the generally accepted guidelines in ROI analysis [31]. The ROIs are shown in Figure 2. The peaks were located in six areas: the left TPJ (Montreal Neurological Institute [MNI]: −52, − 56, 24), right TPJ (MNI: 55, −53, 24), left STS (MNI: −59, −26, −9), right STS (MNI: 59, −18, − 17), MPFC (MNI: 1, 58, 19), and the precuneus (MNI: −3, −56, 37). The radius of the ROI spheres was 10 mm, corresponding to the approximate volume (4,000 mm^3^) of the largest clusters (TPJ and MPFC) reported in the meta-analysis study used here to define the ROIs [18]. The same sphere radius was used for all ROIs.

**Figure 2.**
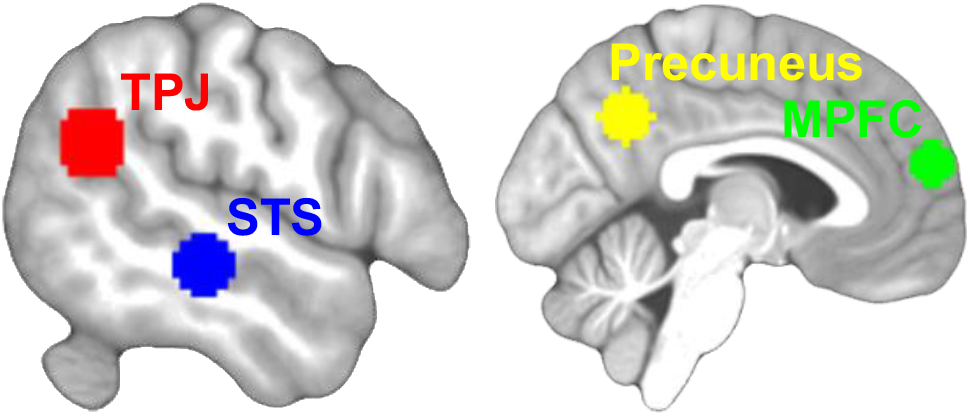
Regions of interest (ROIs). Six ROIs were defined based on peaks reported in an activation likelihood estimation meta-analysis of 16 fMRI studies involving theory-of-mind reasoning [18]. The ROIs consisted of 10-mm-radius spheres centered on peaks in the bilateral temporoparietal junction (TPJ) and superior temporal sulcus (STS), and two midline structures: the precuneus and medial prefrontal cortex (MPFC). Here, the TPJ and STS ROIs on the left side are shown. See Methods for ROI coordinates.

The fMRI data from all participants were analyzed with the Statistical Parametric Mapping software (SPM12) (Wellcome Department of Cognitive Neurology, London, UK) [45]. We first used a conventional general linear model (GLM) to estimate regression beta coefficients for each individual trial (i.e., 256 regressors), focusing on the phase of each trial over 1.5 s immediately after the question mark appeared (the time window in which the subjects were allowed to judge and make a response). A regressor of no interest modeled the initial 4 s of the trial across all conditions. Each regressor was modeled with a boxcar function and convolved with the standard SPM12 hemodynamic response function. In addition, 8 run-specific regressors controlling for baseline differences between runs, and six motion regressors, were included. The trial-wise beta coefficients (i.e., 256 beta maps) were then submitted to subsequent multivariate analyses [46].

The MVPA was carried out using The Decoding Toolbox (TDT) version 3.999 [47] for SPM. For each subject and ROI, we used linear support vector machines (SVMs, with the fixed regularization parameter of C = 1) to compute decoding accuracies. To ensure independent training and testing data sets, we used a leave-one-run-out cross-validation approach. For each fold, an SVM was then trained to discriminate activity patterns belonging to the contrasted trial types in seven runs, and then tested on the trials in the left-out run, repeated for all runs, resulting in a run-average decoding accuracy for each ROI and subject.

For statistical inference, the true group mean decoding accuracy was compared to a null distribution of group mean accuracies obtained from permutation testing. The same MVPA was repeated within each subject and ROI using permuted condition labels (1000 iterations). A p value was computed as (1+the number of permuted group accuracy values > true value)/(1+the total number of permutations). To control for multiple comparisons across the six ROIs, we used the false discovery rate (FDR) correction [48]. In addition, we also computed a bootstrap distribution around the true group mean accuracy by resampling individual-subject mean accuracies with replacement (1000 iterations), from which a 95% confidence interval (CI) was derived [49].

Beyond the targeted hypotheses of this study concerning the six ROIs, we also used a whole-brain searchlight analysis [50] to test for possible areas of decoding outside the ROIs. The searchlight analysis is conceptually different from the ROI analysis. It is not targeted to specific brain areas on the basis of predictions, and therefore is more statistically conservative because of brain-wide multiple comparisons correction. In general, one would not expect the searchlight analysis to align perfectly with the ROI analysis. It is possible to obtain significant results in the ROI analysis that do not appear in the searchlight analysis. Instead, the searchlight analysis is useful for revealing clusters of strong decoding in areas that were not anticipated by hypothesis.

For the searchlight analysis, first, the brain was partitioned into overlapping voxel clusters of spherical shape (10-mm radius). In each of these clusters, a decoding accuracy was computed using the same model input, SVM parameters, and procedures as described for the ROI analysis. For each contrast between two trial types, this process resulted in a decoding accuracy map for each subject, in which the value of each voxel represents the average proportion of correctly classified trials relative to chance level (50%) based on the 10 mm sphere of tissue surrounding that voxel. The subject-wise decoding maps were then smoothed using a 3-mm full-width-half-maximum Gaussian kernel, and entered into a second-level analysis using SPM12. In that analysis, for statistical inference, we employed a cluster-level, whole-brain approach to find clusters that passed the threshold of p < 0.05, corrected for brain-wide multiple comparisons using the family-wise error rate correction as implemented by SPM12.

### Univariate analysis

We subjected the data to univariate analyses to control for potential univariate effects that could contribute to classifier performance in the MVPA. The preprocessed data was smoothed using a 6-mm full-width-half-maximum Gaussian kernel. In the first-level analysis, we modeled the data using the same approach as described above for the MVPA, but defined one regressor per experimental condition (as opposed to one regressor per trial). We then defined linear contrasts in the GLM, and the contrast images from all subjects were entered into a random effects group analysis. For statistical inference, we searched for clusters that passed the threshold of p < 0.05, corrected for multiple comparisons either within each of the six ROIs, or using the whole brain as search space, using the familywise error rate correction as implemented by SPM12.

### Data availability

The data that support the findings of this study are available at https://figshare.com/s/f83f184793f8be13f37f.

## Results

### Task performance

Subjects performed the task at high levels of accuracy, suggesting that they understood the instructions and attributed beliefs to the cartoon heads as intended. (Overall accuracy, 94.8%; for BS trials, 91.6%; BnS trials, 95.2%; nBS trials, 96.0%; nBnS trials, 96.2%; overall lateny = 1006 ms; mean latency for BS trials = 1007 ms, SEM = 29; for BnS trials = 977 ms, SEM = 31; for nBS trials = 1035 ms, SEM = 22; for nBnS trials = 1005 ms, SEM = 28.)

### ROI analysis, Hypothesis 1

Figure 3 and Table 1 show the results of the ROI analysis for hypothesis 1, in which false belief trials (BS) were contrasted with the corresponding true belief trials (BnS). In each panel, the red line shows the accuracy of the MVPA analysis in decoding whether a BS or BnS trial occurred, compared to a chance level of 50%. The histogram shows the null distribution of decoding accuracies based on permutation testing with shuffled conditions labels (1000 iterations). None of the six ROIs showed any significant decoding (the decoding accuracy was not significantly different from the chance level of 50%; see Table 1 for p values and for 95% confidence intervals). We therefore did not find any evidence of a difference between processing false belief and true belief trials, within the six defined ROIs in known ToM cortical areas. Thus, hypothesis 1 was not supported by the analysis within the six ROIs.

**Table 1.** Decoding false belief from true belief trials (BS versus BnS). For definition of ROIs, see Figure 2. Mean decoding accuracy (%), 95% confidence interval (based on bootstrap distribution), and p value (based on permutation testing, uncorrected for multiple comparisons) are shown for each of the six ROIs. The * indicates significant p values that survived correction for multiple comparisons across all six ROIs (FDR-corrected p < 0.05).

|  | **L TPJ** | **R TPJ** | **L STS** | **R STS** | **MPFC** | **Precuneus** |
| --- | --- | --- | --- | --- | --- | --- |
| Decoding accuracy | 48.6% | 50.6% | 50.1% | 49.7% | 49.4% | 51.4% |
| 95% CI | 47.5 to 50.3 | 49.0 to 52.3 | 48.4 to 51.6 | 47.9 to 51.8 | 47.3 to 51.3 | 49.8 to 52.9 |
| P value | 0.936 | 0.260 | 0.420 | 0.683 | 0.776 | 0.0530 |

**Figure 3.**
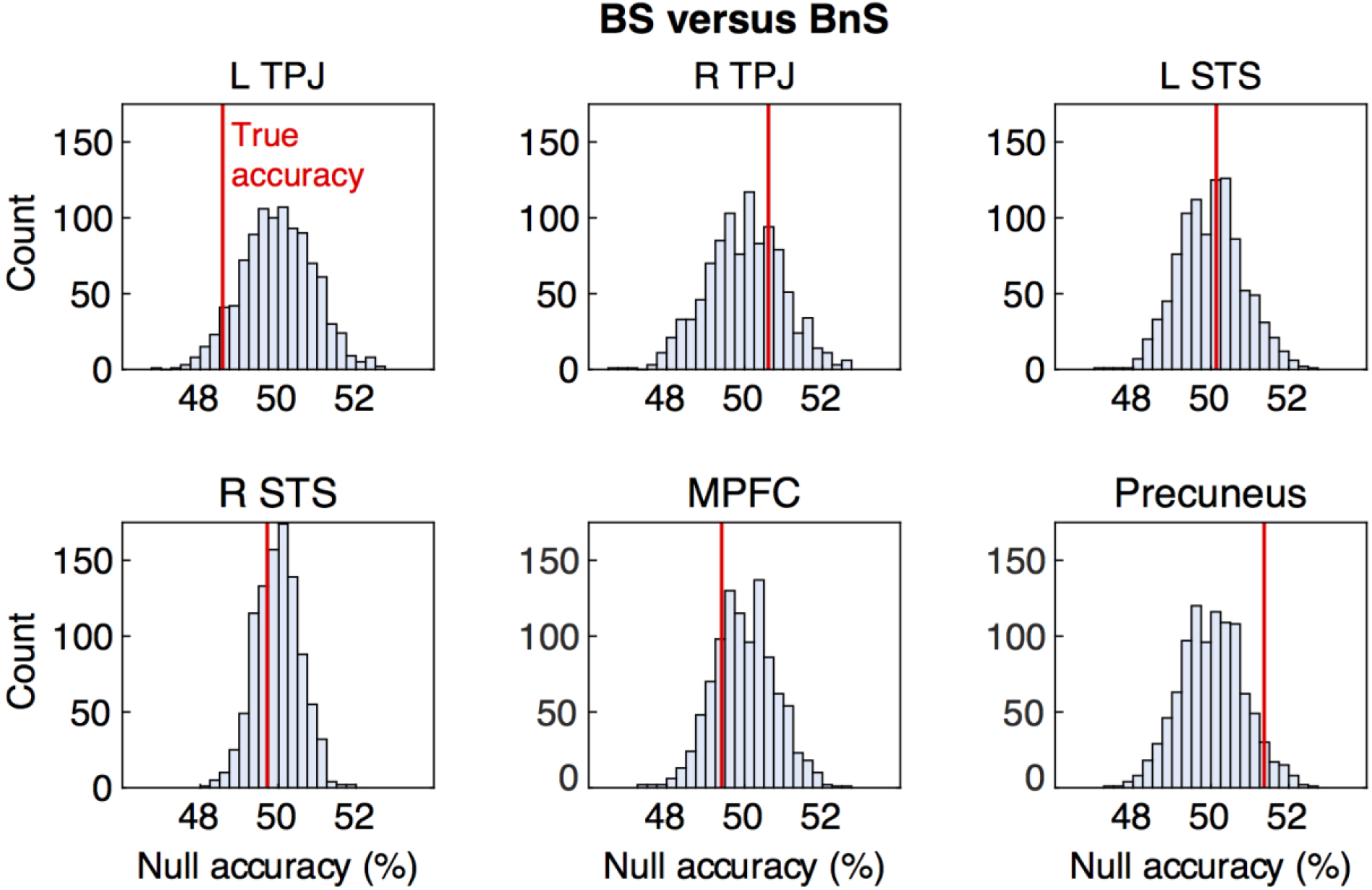
Decoding false belief from true belief trials. False belief trials were represented by the BS condition and matching true belief trials by the BnS condition. For definition of the six ROIs, see Figure 2. Each panel shows the results for one ROI. In each panel, the histogram shows the null distribution of decoding accuracies based on permutation testing with shuffled conditions labels (chance level = 50%). The red vertical line shows the accuracy of the classifier when it was trained and tested using the real (unshuffled) conditions labels. Significance threshold (p<0.05) based on permutation testing, corrected for multiple comparisons across six ROIs using FDR. None of the ROIs showed significant decoding that distinguished false from true belief trials.

### ROI analysis, Hypothesis 2

Figure 4 and Table 2 show the results for the ROI analysis for hypothesis 2, in which nBS trials were contrasted with nBnS trials. According to hypothesis 2, at least some of the ROIs should show significant decoding for this contrast, indicating sensitivity to switched versus nonswitched trials in the specific case in which the head’s vision is unblocked and able to “see” whether the ball has been switched or not. The results show a significant decoding for nBS versus nBnS trials in the right TPJ (the decoding accuracy was significantly above the chance level of 50%). This decoding result was significant even when corrected for multiple comparisons across the six defined ROIs (p = 0.004 uncorrected; p = 0.024 corrected using FDR; see also 95% confidence intervals in Table 2).

**Table 2.** Decoding trials in which the cartoon “saw” a switch from trials in which the cartoon “saw” no switch (nBS versus nBnS). For definition of ROIs, see Figure 2. Mean decoding accuracy (%), 95% confidence interval (based on bootstrap distribution), and p value (based on permutation testing, uncorrected for multiple comparisons) are shown for each of the six ROIs. The * indicates significant p values that survived correction for multiple comparisons across all six ROIs (FDR-corrected p < 0.05).

|  | **L TPJ** | **R TPJ** | **L STS** | **R STS** | **MPFC** | **Precuneus** |
| --- | --- | --- | --- | --- | --- | --- |
| Decoding accuracy | 49.8% | 52.8% | 49.4% | 50.2% | 49.7% | 50.4% |
| 95% CI | 47.7 to 52.1 | 50.9 to 54.6 | 47.7 to 50.9 | 48.2 to 52.0 | 48.4 to 51.6 | 48.6 to 52.1 |
| P value | 0.518 | 0.004* | 0.725 | 0.354 | 0.631 | 0.3290 |

**Figure 4.**
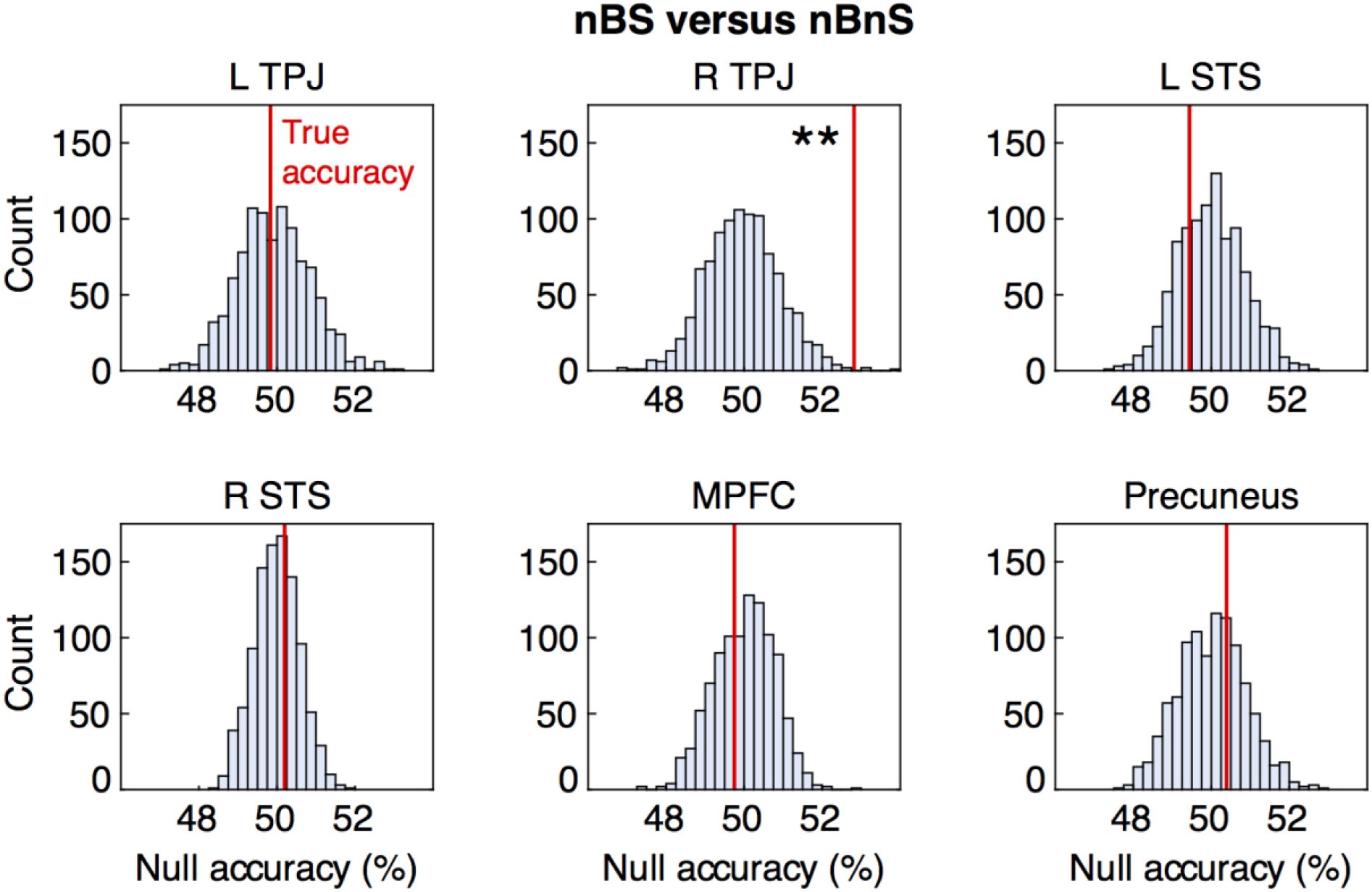
Decoding trials in which the cartoon “saw” a switch occur from trials in which the cartoon “saw” that no switch occurred. Trials in which the cartoon “saw” a switch were represented by the nBS condition, and trials in which the cartoon “saw” no switch were represented by the nBnS condition. For definition of the six ROIs, see Figure 2. Each panel shows the results for one ROI. In each panel, the histogram shows the null distribution of decoding accuracies based on permutation testing with shuffled conditions labels (chance level = 50%). The red vertical line shows the accuracy of the classifier when it was trained and tested using the real (unshuffled) conditions labels. A decoding accuracy significantly greater than chance is indicated by * (p<0.05), based on permutation testing, corrected for multiple comparisons across six ROIs using FDR. The right TPJ showed significant decoding (p corrected = 0.024).

Finally, we compared the strength of the decoding obtained in the hypothesis 2 test to the strength of the decoding obtained in the hypothesis 1 test. The decoding in the right TPJ for nBS versus nBnS trials (hypothesis 2 test) was significantly greater than the decoding in the right TPJ for BS versus BnS trials (hypothesis 1 test) (52.8% versus 50.6%; p = 0.0432, permutation testing with 10,000 iterations). Thus, hypothesis 2 was supported by the present data and hypothesis 1 was not. When subjects thought that the head could see the switch take place, then the right TPJ was affected by the switch (significant decoding for nBS versus nBnS); when subjects thought that the head could not see the switch take place, then none of the ToM ROIs was significantly affected by the switch (no significant decoding for BS versus BnS).

### Searchlight analysis

As a further exploration beyond the targeted hypotheses of this study, we used a whole-brain searchlight analysis [50] to test for possible areas of decoding outside the ROIs. Because the searchlight analysis does not test strong a priori hypotheses and requires statistical correction across the full brain, it is much less sensitive. The searchlight comparison between BS and BnS trials revealed no significant areas of decoding at the brain-wide level; likewise, the searchlight comparison between nBS and nBnS trials revealed no significant areas of decoding at the brain-wide level.

### Univariate analysis

To control for potential univariate effects that could drive classifier performance in the decoding analyses, we examined the bi-directional contrasts for the BS-versus-BnS and the nBS-versus-nBnS comparisons (i.e., BS > BnS, BnS > BS, nBS > nBnS, and nBnS > nBS). None of the contrasts revealed significant activity, neither within the ROIs nor at the whole-brain level.

The same result was found for all six possible specific comparisons (12 contrasts) between individual conditions, the two main effects (4 contrasts), and the interaction (2 contrasts). The absence of any univariate effect within the ROIs, or anywhere else in the brain, confirm that the stimuli were well matched. These findings are compatible with previous studies [51] that demonstrated the superiority of pattern-sensitive multivariate analyses compared to conventional univariate approaches for detecting differences in activity between conditions with highly similar macroscopic characteristics.

## Discussion

The present experiment used fMRI to measure brain activity during a pictorial, reaction-time, ToM task that incorporated both false belief and true belief trials. We used the task to test two specific hypotheses. In hypothesis 1, brain areas in a previously described ToM cortical network should respond differently to false belief trials than to true belief trials (contrasting BS and BnS trials). This hypothesis was not supported by the data. We found no significant decoding of false belief versus true belief trials in any of the six ROIs that we tested. In hypothesis 2, the ToM cortical network should not respond differently to BS and BnS trials, because the cartoon head had its vision blocked in both conditions – subjects knew that it could not “see” the ball switch – and thus, by implication, the mental state of the head should have been unaffected by whether the ball switched or not. The ToM brain areas, building a model of the mental state of the agent, therefore should not react differently in those two conditions.

Instead, only when the head was unblocked, and by implication could see whether the ball switched or not, should the ToM network react differently to the switch and nonswitch conditions. Hypothesis 2 therefore predicted significant decoding in the nBS versus nBnS comparison. This hypothesis was supported by the data. The right TPJ showed decoding significantly above chance for the nBS versus nBnS comparison.

A possible alternative explanation of the present results is that the right TPJ reacted to the difference between switched and nonswitched trials because of a processing cost, or other general effect on processing, caused by the subjects seeing the ball switch from one box to another. The average percent correct was indeed slightly lower (by an average of 1.9%), and the average latency higher (by 30 ms), in switched conditions than in nonswitched conditions (see **Results**, Task performance), suggesting a processing cost on switched trials. However, this difference in performance is unlikely to explain the TPJ result. The difference was larger for BS versus BnS trials, than it was for nBS versus nBnS trials. For the BS versus BnS trials, we found no significant signal in the right TPJ; and for the nBS versus nBnS trials, we did find a significant signal in the right TPJ; and the two comparisons were significantly different from each other. If the right TPJ activity reflected a difference between switched and nonswitched trials, it was evidently not a general effect, but only occurred when the subjects thought that the cartoon character could see the switch take place. We suggest, therefore, that our interpretation in terms of modeling the mind states of others is the most likely.

The present results might also help to explain the somewhat mixed results of previous studies that compared false belief and true belief conditions [23,25,28,29]. On the one hand, false belief conditions may require more cognitive complexity or effort on the part of the subject. For that reason, one might hypothesize that the ToM cortical network should be more active in false belief trials than in true belief trials. On the other hand, the implied mental state of the agent in question is not necessarily different in false versus true belief trials. Thus, by modeling the same mental state, the ToM cortical network might respond in the same way to false and true belief trials. Comparing false and true belief trials, therefore, may be a less incisive test of the ToM network than comparing two different mental states attributed to an agent. On the basis of the current data, we cannot rule out the possibility that brain areas in the ToM network, such as the right TPJ, react to the distinction between false and true beliefs in others. We did not find that signal, but perhaps our paradigm was not sensitive enough to detect it. For the sake of caution, therefore, what we can infer from our results is that in the particular task that we used, when analyzing brain activity at the moment of a reaction-time decision, the right TPJ, a central part of the ToM network, appears to place greater emphasis on reconstructing the internal mental experience of another agent than it does on the outside perspective or knowledge that allows the subject to evaluate whether that agent has a false or true belief.

The present experimental design allowed for a variety of exploratory comparisons between task conditions, beyond the hypothesis-driven BS-versus-BnS comparison and nBS-versus-nBnS comparison reported above. The **Supplementary Information** provides the results of the full set of comparisons between task conditions.

## Acknowledgements

Supported by the Princeton Neuroscience Institute Innovation Fund and Princeton Program in Cognitive Science. Arvid Guterstam was supported by the Wenner-Gren Foundation, the Swedish Brain Foundation, and the Promobilia Foundation. We thank Jemma Bio-Barrick and Argos Wilterson for their encouragement.

## Supplementary Information

In the main text, we describe using MVPA to contrast BS to BnS trials, and nBS to nBnS trials. These two contrasts were motivated by specific hypotheses about the cortical ToM network. Here, for completeness, we report the results of eight contrasts. These contrasts include the two main factors: all blocked trials versus all nonblocked trials, and all switched trials versus all nonswitched trials. The contrasts also include all possible pairs among the four specific conditions: BS versus BnS, BS versus nBS, BS versus nBnS, BnS versus nBS, BnS versus nBnS, and nBS versus nBnS. For each contrast, we performed the MVPA analysis on the six defined ROIs. Figure S1 and Table S1 show the results. Only two analyses show significant decoding. The blocked versus nonblocked contrast resulted in significant decoding in the precuneus (p uncorrected = 0.006; p corrected for 6 ROI = 0.036). The nBS versus nBnS contrast resulted in significant decoding in the rTPJ, as reported in the main text (p uncorrected = 0.004; p corrected for 6 ROI = 0.024).

For each of the eight contrasts, we also performed a brain-wide, searchlight MVPA analysis. None of these analyses revealed any clusters of significant decoding, corrected for multiple comparisons across the brain.

**Caption for Table S1.** Eight different contrasts, each performed within the six ROIs. Mean decoding accuracy (%), 95% confidence interval (based on bootstrap distribution), and p value (based on permutation testing, uncorrected for multiple comparisons) are shown for each of the six ROIs. The * indicates significant p values that survived correction for multiple comparisons across all six ROIs (FDR-corrected p < 0.05).

|  |  | **L TPJ** | **R TPJ** | **L STS** | **R STS** | **MPFC** | **Precuneus** |
| --- | --- | --- | --- | --- | --- | --- | --- |
| **Blocked**<br>**versus**<br>**nonBlocked** | Decoding<br>accuracy (%) | 51.4 | 50.4 | 50.7 | 50.0 | 50.5 | 51.5 |
|  | 95% CI | 49.7 to 52.8 | 49.1 to 51.6 | 49.7 to 51.9 | 48.5 to 51.4 | 49.6 to 51.4 | 50.2 to 53.2 |
|  | P value | 0.021 | 0.269 | 0.115 | 0.448 | 0.173 | 0.006* |
| **Switched**<br>**versus**<br>**NonSwitched** | Decoding<br>accuracy (%) | 50.2 | 50.4 | 50.7 | 49.5 | 49.3 | 49.9 |
|  | 95% CI | 48.9 to 51.6 | 49.0 to 51.6 | 49.2 to 52.0 | 48.6 to 50.6 | 48.1 to 50.5 | 48.8 to 51.1 |
|  | P value | 0.373 | 0.275 | 0.126 | 0.880 | 0.912 | 0.588 |
| **BS**<br>**versus**<br>**BnS** | Decoding<br>accuracy (%) | 48.6 | 50.6 | 50.1 | 49.7 | 49.4 | 51.4 |
|  | 95% CI | 47.5 to 50.3 | 49.0 to 52.3 | 48.4 to 51.6 | 47.9 to 51.8 | 47.3 to 51.3 | 49.8 to 52.9 |
|  | P value | 0.936 | 0.260 | 0.420 | 0.683 | 0.776 | 0.053 |
| **BS**<br>**versus**<br>**nBS** | Decoding<br>accuracy (%) | 51.1 | 50.7 | 50.9 | 51.2 | 48.7 | 50.4 |
|  | 95% CI | 49.3 to 53.1 | 48.7 to 52.5 | 48.9 to 53.0 | 49.4 to 52.7 | 46.9 to 50.6 | 48.6 to 51.9 |
|  | P value | 0.137 | 0.243 | 0.163 | 0.021 | 0.960 | 0.305 |
| **BS**<br>**versus**<br>**nBnS** | Decoding<br>accuracy (%) | 50.7 | 50.4 | 50.7 | 50.4 | 50.1 | 50.0 |
|  | 95% CI | 49.2 to 52.5 | 48.6 to 52.3 | 49.1 to 52.4 | 49.0 to 52.1 | 48.3 to 52.0 | 48.4 to 51.6 |
|  | P value | 0.218 | 0.318 | 0.199 | 0.252 | 0.409 | 0.462 |
| **BnS**<br>**versus**<br>**nBS** | Decoding<br>accuracy (%) | 50.9 | 49.6 | 51.2 | 50.2 | 49.9 | 49.9 |
|  | 95% CI | 48.7 to 53.0 | 47.3 to 51.5 | 49.1 to 53.2 | 48.2 to 52.1 | 48.0 to 52.2 | 47.9 to 51.9 |
|  | P value | 0.183 | 0.678 | 0.081 | 0.320 | 0.515 | 0.529 |
| **BnS**<br>**versus**<br>**nBnS** | Decoding<br>accuracy (%) | 51.4 | 49.8 | 50.4 | 48.8 | 50.1 | 49.7 |
|  | 95% CI | 49.1 to 53.5 | 48.4 to 51.4 | 48.2 to 52.7 | 46.7 to 50.6 | 48.2 to 51.8 | 48.0 to 51.7 |
|  | P value | 0.071 | 0.583 | 0.329 | 0.976 | 0.459 | 0.633 |
| **nBS**<br>**versus**<br>**nBnS** | Decoding<br>accuracy (%) | 49.8 | 52.8 | 49.4 | 50.2 | 49.7 | 50.4 |
|  | 95% CI | 47.7 to 52.1 | 50.9 to 54.6 | 47.7 to 50.9 | 48.2 to 52.0 | 48.4 to 51.6 | 48.6 to 52.1 |
|  | P value | 0.518 | 0.004* | 0.725 | 0.354 | 0.631 | 0.329 |

**Figure S1.**
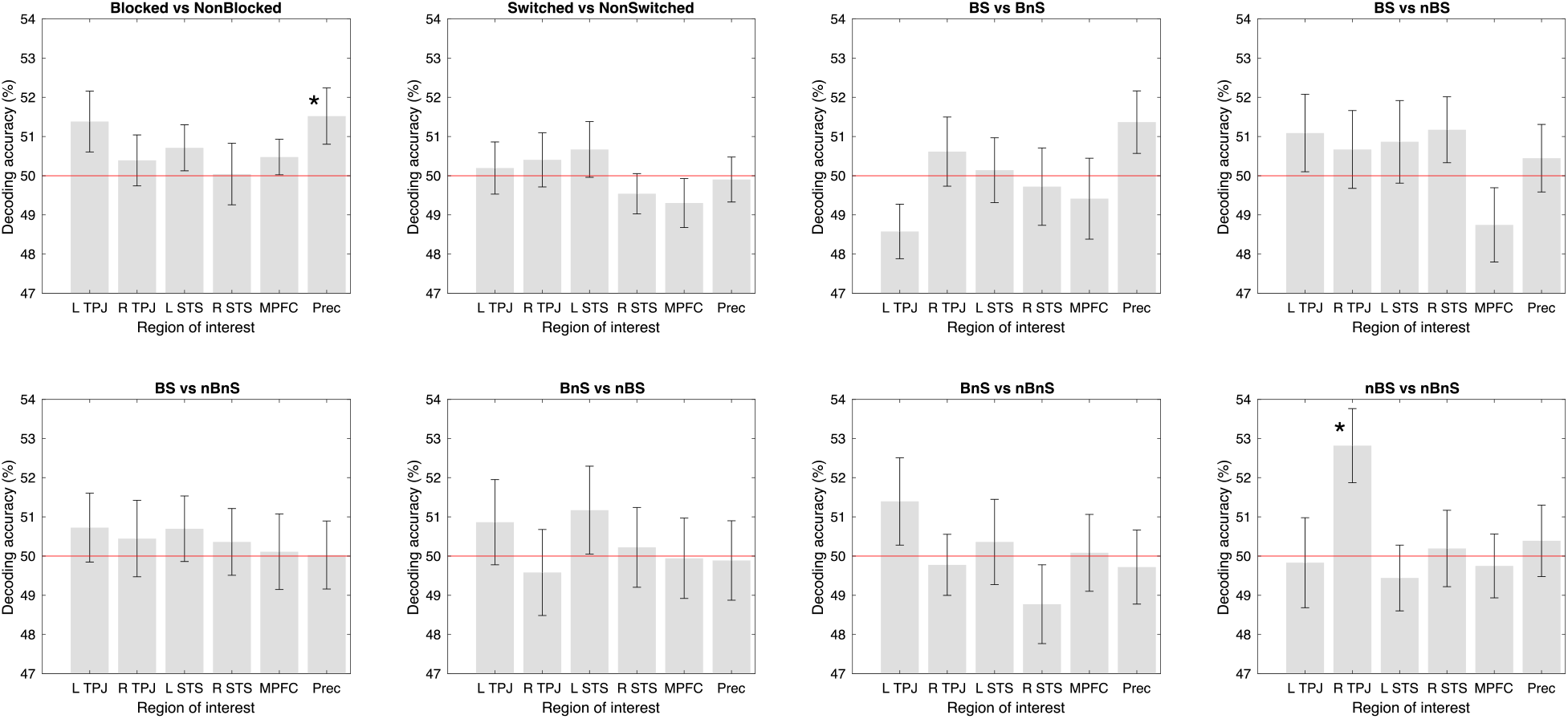
Eight different contrasts, each performed within the six ROIs. Each bar shows mean decoding accuracy. Error bars show SEM. Red horizontal line indicates chance level decoding. Significance indicated by * (p<0.05) based on permutation testing, corrected for multiple comparisons across the six ROIs using FDR.

## Notes

https://figshare.com/s/f83f184793f8be13f37f

### Competing Interest Statement

The authors have declared no competing interest.

